# Distinct Classes of Lamin-Associated Domains are Defined by Differential Patterns of Repressive Histone Methylation

**DOI:** 10.1101/2024.12.20.629719

**Authors:** Caden J. Martin, Prabakaran Nagarajan, Elizabeth A. Oser, Liudmila V. Popova, Mark R. Parthun

## Abstract

A large fraction of the genome interacts with the nuclear periphery through lamina-associated domains (LADs), repressive regions which play an important role in genome organization and gene regulation across development. Despite much work, LAD structure and regulation are not fully understood, and a mounting number of studies have identified numerous genetic and epigenetic differences within LADs, demonstrating they are not a uniform group. Here we profile Lamin B1, HP1β, H3K9me3, H3K9me2, H3K27me3, H3K14ac, H3K27ac, and H3K9ac in MEF cell lines derived from the same mouse colony and cluster LADs based on the abundance and distribution of these features across LADs. We find that LADs fall into 3 groups, each enriched in a unique set of histone modifications and genomic features. Each group is defined by a different heterochromatin modification (H3K9me3, H3K9me2, or H3K27me3), suggesting that all three of these marks play important roles in regulation of LAD chromatin and potentially of lamina association. We also discover unique features of LAD borders, including a LAD border-specific enrichment of H3K14ac. These results reveal important distinctions between LADs and highlight the rich diversity and complexity in LAD structure and regulatory mechanisms.

## Introduction

Histone post-translational modifications (PTMs) help shape the 3-dimensional organization of the genome. Combinatorial patterns of histone PTMs distinguish different states of chromatin, interact with transcription factors and chromatin binding proteins to mediate transcriptional activation and repression, and help regulate the conformation and physical properties of chromatin(Janssen and Lorincz 2022; Macrae et al. 2023). Histone PTMs also play a crucial role in transmitting the memory of chromatin state and cell identity through cell division(Escobar et al. 2021).

Most actively transcribed genes are found in euchromatin, which is relatively decondensed, enriched in histone acetylation, transcription factor binding, and RNA polymerase II. Euchromatin largely localizes to the nuclear interior. Heterochromatin is transcriptionally repressive and highly compacted. There are two fundamentally distinct forms of heterochromatin. Constitutive heterochromatin is enriched in H3K9 di- and tri-methylation (H3K9me2/3) and the H3K9me2/3- binding protein HP1, depleted in histone acetylation, and generally localizes to the nuclear periphery or chromocenters(Allshire and Madhani 2018; Grewal 2023). Facultative heterochromatin is likewise depleted in histone acetylation but is enriched in H3K27 trimethylation (H3K27me3) rather than H3K9 methylation. Both forms of heterochromatin contribute to gene repression, though likely through different mechanisms. Facultative heterochromatin tends to silence genes in a developmental or cell-type specific manner(Kim and Kingston 2022). Constitutive heterochromatin blocks transcription of constitutively silent regions like centromeres, telomeres, and repetitive elements in a non-cell-type-specific manner, although it has recently emerged that regions with features characteristic of constitutive heterochromatin can also function in cell type-specific gene regulation(Nicetto et al. 2019; Nicetto and Zaret 2019). Constitutive heterochromatin also likely functions independent of its role in transcriptional repression to regulate genome architecture and stability(Falk et al. 2019).

Lamin-associated domains (LADs) are large, heterochromatic regions of the genome that interact with the nuclear lamina at the inner nuclear membrane, an environment which generally promotes gene silencing. Some LADs are conserved across development, while others interact with the lamina only in particular cell types, leading to the subdivision of LADs into constitutive LADs (cLADs) and facultative LADs (fLADs) (not to be confused with constitutive and facultative heterochromatin)(Peric-Hupkes et al. 2010; Shah et al. 2023). Multiple mechanisms mediate the physical interaction of LADs with the nuclear periphery(Manzo et al. 2022).

LADs are classically characterized by marks of constitutive heterochromatin. H3K9me2 and H3K9me3 are enriched along LAD bodies and are the primary modifications involved in tethering to the nuclear periphery(Bian et al. 2013; van Steensel and Belmont 2017). Several studies have emphasized the significance of H3K9me2 in promoting chromatin-lamina interactions, showing that H3K9me2 is enriched in LADs and around the nuclear periphery and that depletion or overexpression of the H3K9me2-methyltransferase G9a causes reduction and enhancement, respectively, of chromatin-lamina interactions(Peric-Hupkes et al. 2010; Kind et al. 2013; Poleshko et al. 2017; Poleshko et al. 2019; Smith et al. 2021). H3K9me3 has also been shown to localize to the nuclear lamina and promote the peripheral localization of chromatin(Bian et al. 2013; Keenan et al. 2024). H3K9me2/3 both recruit Heterochromatin Protein 1 (HP1), an important mediator of gene silencing and gene compaction which helps link chromatin to the periphery through interactions with A and B-type lamins and lamin-associated proteins such as LBR and PRR14(Ye and Worman 1996; Poleshko et al. 2013).

The facultative heterochromatin mark H3K27me3 has been seen to be specifically enriched at LAD borders and some smaller, more gene-rich LADs, though little attention has been paid to the latter category(Guelen et al. 2008; Sadaie et al. 2013; Harr et al. 2015; Tran et al. 2021; Alagna et al. 2023; Gholamalamdari et al. 2024). H3K27me3 peaks at LAD borders are believed to insulate the constitutive heterochromatin inside LADs from euchromatin outside of the LAD(Guelen et al. 2008; Siegenfeld et al. 2022). H3K27me3 may also help regulate the association of LADs with the nuclear lamina, though the extent of its importance is unclear and studies conflict as to whether it promotes or inhibits chromatin-lamina interactions(Harr et al. 2015; Guerreiro and Kind 2019; Siegenfeld et al. 2022).

Multiple factors complicate our understanding of LAD structure and regulation. One is the large degree of redundancy between tethering mechanisms and the diversity in peripheral proteins which can interact with chromatin, many of which are differentially expressed between different tissues and stages of development. Added to this is the variability of LADs themselves. While they are consistently heterochromatic and generally transcriptionally repressive, LADs show mixed abilities to silence genes and gene reporters, which is often dependent on the chromatin context and the promoter itself(Leemans et al. 2019; Alagna et al. 2023). Previous studies have uncovered a lack of uniformity across LADs in their MNase accessibility, linker histone enrichment, lamin B1 occupancy, CTCF dependency, primary lamin interactor (B or A/C), histone PTM enrichment, and more(Lund et al. 2015; Zheng et al. 2015; Kaczmarczyk et al. 2022; Alagna et al. 2023; Shah et al. 2023). While some degree of variation across the genome is expected for any set of genomic features, these studies suggest that LADs cannot be simply characterized as a single group and may fall into subcategories with distinct structures, functions and modes of regulation.

To better understand the structure and regulation of lamina-associated chromatin, we determined the genome-wide localization of Lamin B1 (CUT&RUN), HP1β (ChIP-seq), and 6 different histone modifications (CUT&Tag) across the same set of immortalized mouse embryonic fibroblast (iMEF) cell lines derived from a single mouse colony. We then grouped LADs according to the levels of each feature throughout the LAD. We find that LADs fall into 3 classes which exhibit distinct chromatin profiles, gene enrichments, DNA replication timing and size distributions. These results uncover novel features of LAD structure, highlight underrecognized differences between LADs, and underscore the importance of considering LAD subtype when studying chromatin-lamina interactions.

## Results

### Comparison of LADs derived from a single set of related cell lines

The goal of this study was to obtain a more comprehensive understanding of LAD structure by profiling the patterns of multiple histone PTMs and chromatin associated proteins from the same cell line. We chose to use immortalized MEF cell lines derived from wild type C57/bl6 mice. LADs are defined as regions of the genome which interact with the nuclear lamina. We profiled LADs in our iMEFs by performing CUT&RUN for Lamin B1. We compared our LADs with a previously published map of LADs created in NIH3T3 MEFs using LaminB1-DamID. Overall, DamID indicated that LADs comprise 40.7% of the genome in NIH3T3 cells while CUT&RUN in C57/bl6 iMEFs indicated that LADs cover 48.7% of the genome (Figure 1A). There is extensive overlap between the sets of LADs with 88% of the DamID LADs also identified by CUT&RUN.

**Figure 1:**
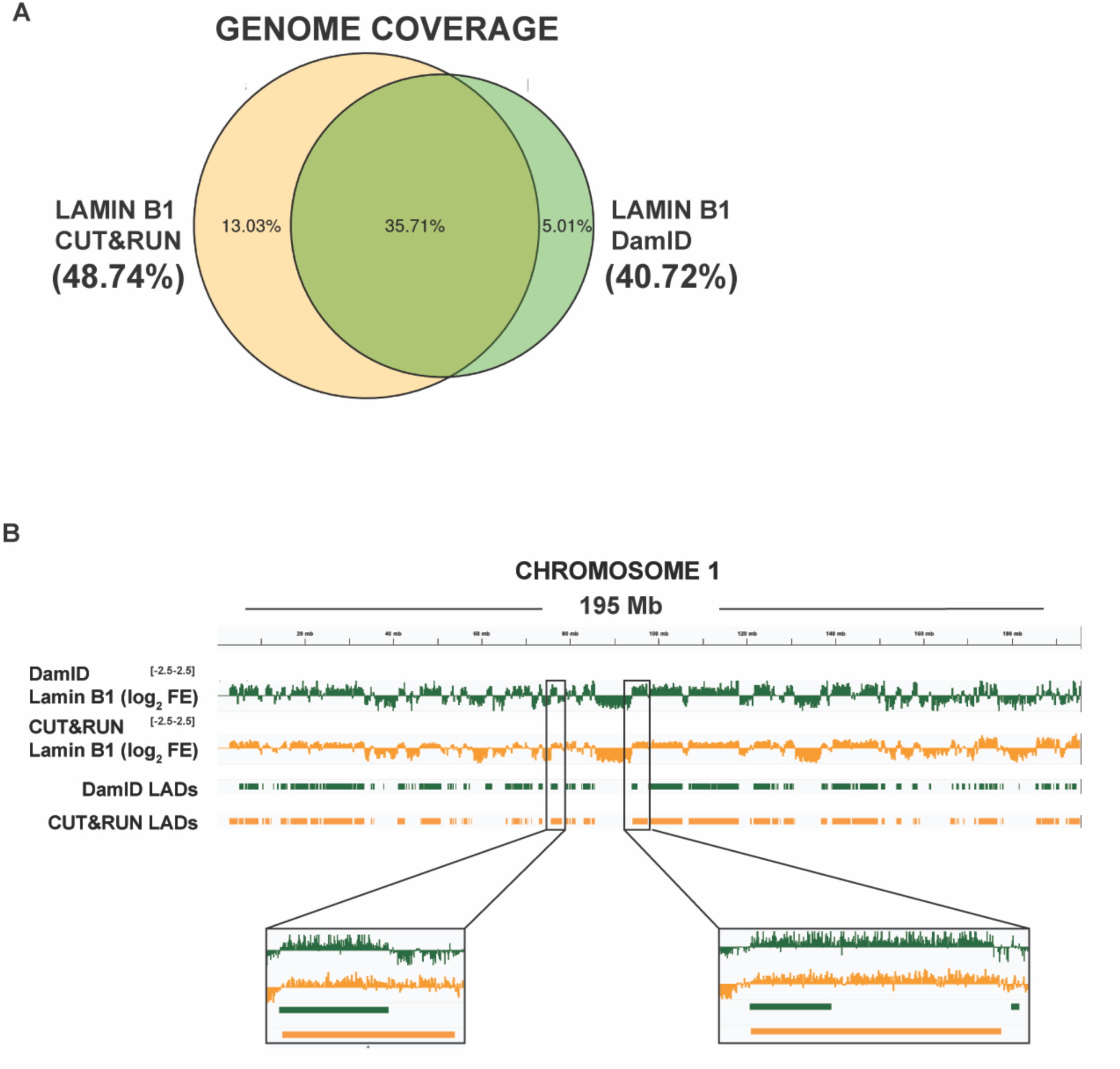
Mapping interaction of chromatin with Lamin B1 in MEFs using CUT&RUN. (A) Fraction of base pairs in the genome covered by our CUT&RUN-generated LAD map alone (yellow), a DamID-generated LAD map in NIH3T3 cells (green), or both. (B) Lamin B1 log2 fold-enrichment over IgG illustrating differences between our CUT&RUN-generated map and the DamID-generated map. Left inset shows differences between maps arising from biological differences between cell lines. Right inset shows differences between maps arising from differences in LAD calling.

The overlap between LADs detected by DamID and CUT&RUN is illustrated by tracks of the log_2_ fold enrichment of lamin B determined by each method (Figure 1B). Much of the difference in LAD detection can be attributed to two factors. The first is biological differences between the two different MEF cell lines. In the lower left inset of Figure 1B, the LAD detected by CUT&RUN is significantly larger due to clear detection of lamin B interaction in the C57/bl6 MEFs that is not present in the NIH3T3 MEFs used for DamID. The second factor is related to the bioinformatic methods used to call LADs. The published dataset used a custom Hidden Markov Model (HMM)-based approach to identify LADs from Lamin B1-DamID data. We also used a HMM to call LADs, but we used a modified version of the HMMt package in R and optimized the parameters for our CUT&RUN data (see Methods for details). For the LAD shown in the lower right inset of Figure 1B, there is clear lamin B1 chromatin association across a large domain. Our method identifies this entire region as a LAD while the DamID method identified a much smaller segment as a LAD. The reverse is also true as there are examples where regions of lamin association were called by the DamID method but not by the CUT&RUN method (data not shown). For subsequent analyses, we use the LAD definitions determined by CUT&RUN in the C57/bl6 iMEFs. When we repeat these analyses using previous LAD set from NIH3T3 cells, we obtain largely similar results (data not shown).

### LAD Borders Show Enrichment of Multiple Histone Modifications

To determine the structure of chromatin across LADs, we used CUT&Tag to map a variety of histone PTMs, including the euchromatin marks H3K9ac, H3K14ac, and H3K27ac, facultative heterochromatin mark H3K27me3, and constitutive heterochromatin marks H3K9me2 and H3K9me3. We also used ChIP-seq to profile the H3K9 methylation binding protein Heterochromatin Protein 1 beta (HP1β).

Scaling all LADs to a uniform size, we plotted the abundance of each feature across the LAD body and around the LAD border (Figure 2A). As expected, we saw broad domains of HP1, lamin B1, and the constitutive heterochromatin mark H3K9me3 across the interior of the LADs. We observed a different pattern for the other constitutive heterochromatin mark, H3K9me2. H3K9me2 is present as a sharp peak just inside of LADs that decreases to an intermediate level across the interior of LADs. The facultative heterochromatin mark H3K27me3 was also present as a peak just inside of LAD borders that then sharply decreased across the LAD interiors to levels at or below the surrounding chromatin, as previously described.^17–19^

**Figure 2:**
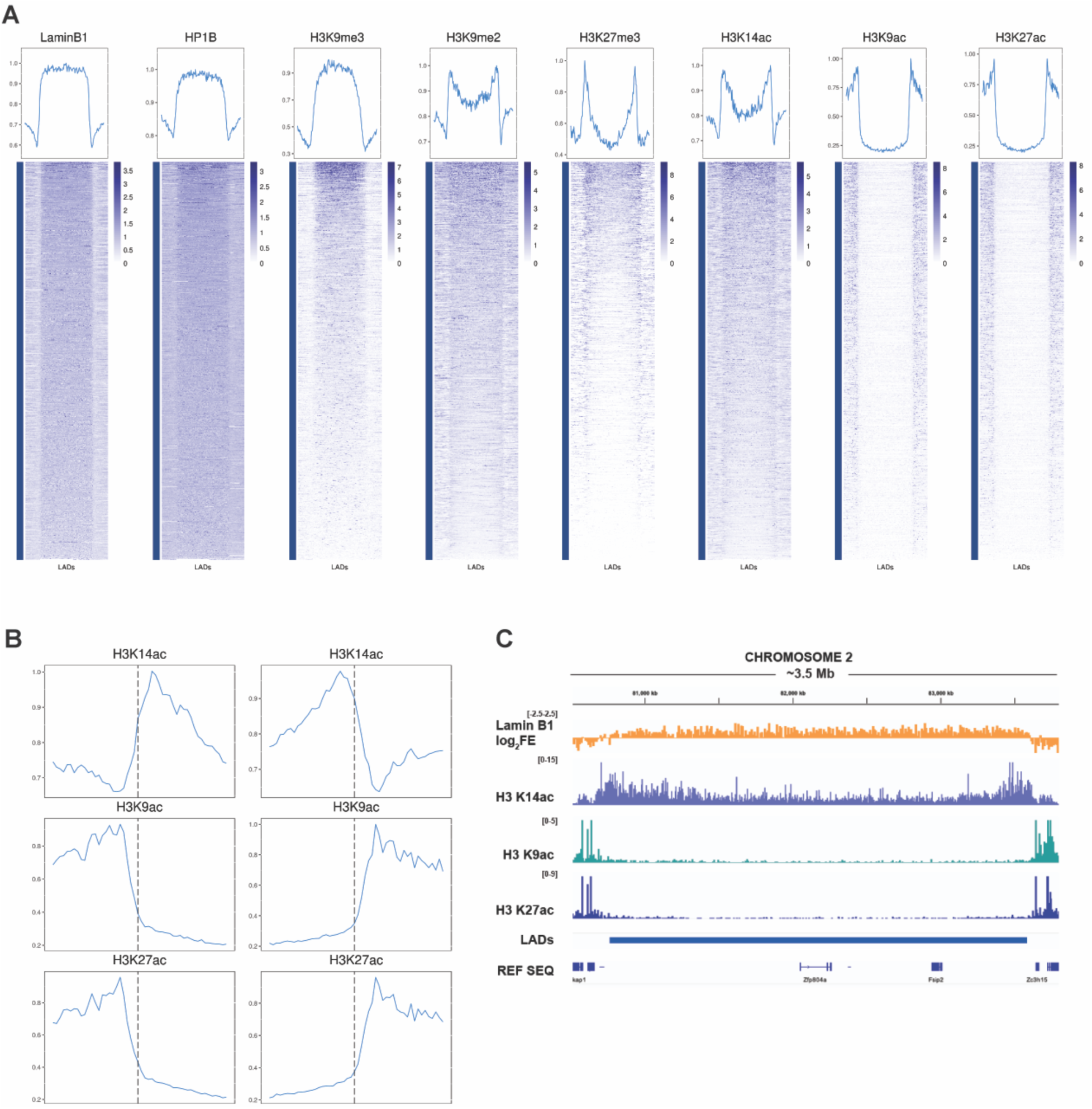
Chromatin profiles across LADs and at LAD borders. (A) Heatmaps of feature abundance across LADs scaled to the same size. Profiles above represent the average signal across all LADs and extend 100 kb beyond each LAD border. (B) Profiles of H3K14ac, H3K9ac, and H3K27ac around LAD borders. Boxes show a 1 Mb window centered around the LAD borders (vertical dotted line). Note that the profiles extend 500 kb inside the LAD border, longer than the full length of some LADs, and thus include the opposite LAD border. This results in the slight increase in H3K9/27ac in the middle of the LAD. (C) IGV Browser view of Lamin B1, H3K14ac, H3K9ac, and H3K27ac across an example LAD.

In addition to the presence of H3K27me3 and H3K9me2 peaks just inside LAD borders, we also observed a complex pattern of histone acetylation around LAD borders. H3K9ac and H3K27ac were essentially depleted throughout the interior of LADs then abruptly increased outside LAD borders (Figure 2B and 2C). H3K14ac showed a much different distribution. H3K14ac formed a broad peak inside of the LAD border, then gradually decreases across the interior of the LAD (Figure 2B and 2C). The presence of significant levels of H3K14ac throughout the interior of LADs indicates that these regions are not entirely devoid of histone acetylation but are permissive to specific sites of acetylation. Strikingly, the locations of the H3K14ac and H3K27me3 peaks did not coincide at LAD borders. H3K27me3 reaches its maximum level exactly at the LAD border, while H3K14ac reaches its maximum a few kilobases to tens of kilobases inside the LAD. Thus, these two modifications may function together to form a barrier insulating LAD chromatin from the surrounding euchromatin.

To confirm that these unanticipated enrichments of H3K9me2 at LAD borders could not be attributed to non-specific binding or unsuitability of the antibody for CUT&Tag, we used CUT&RUN to profile H3K9me2 using the same antibody and using an alternative antibody. We included Epicypher’s K-MatStat Panel of barcoded post-translationally modified nucleosomes to verify antibody specificity for H3K9me2. Results from the K-MetStat panel showed that our H3K9me2 antibody was indeed specific for H3K9me2 (Figure S1A), and visualization of the CUT&RUN data showed highly similar distributions between the two different antibodies and between CUT&RUN and CUT&Tag (Figure S1B). Similarly, we profiled H3K14ac using CUT&RUN with an alternative antibody and determined that the genomic distribution of H3K14ac was unchanged between CUT&Tag data generated with the original antibody and CUT&RUN data generated by the second antibody (Figure S1C).

These results highlight the underappreciated complexity of LAD borders, where multiple complementary mechanisms may be at play to insulate different categories of chromatin and regulate the interaction of LADs with the nuclear periphery.

### LADs Cluster into Distinct Subtypes

While LADs are consistently heterochromatic and contribute to gene silencing, numerous prior studies have found variation among LADs in both their physical and chemical properties. This suggests the possibility that the chromatin profiles created across all LADs in aggregate (Figure 2) mask nuance in the chromatin profiles of individual LADs or groups of LADs. To address this question, we clustered LADs by scaling them all to a uniform size and dividing the chromatin signal into 24 bins across each LAD, then performing k-means clustering on the resulting matrix. Elbow and Silhouette plots suggested 4 clusters to be optimal (Figure 3A, Figure S2A). However, we found that one of these clusters consisted of a small number of LADs. PCA analysis (Figure 3B) indicated that the 3 clusters form distinct groups and that the cluster containing the small number of LADs could be grouped with cluster 2 LADs, which are most similar to it (see the gold outlying point at the top of Figure 3B). Similar results were obtained using hierarchical clustering (Figure S2B/C). Thus, we ran our analysis using k-means clustering with 3 clusters. Only LADs on autosomal chromosomes were included due to lower quality Lamin B1 mapping on sex chromosomes and sex differences between the mice from which the replicate cell lines were derived. The resulting three clusters contained 409, 407, and 312 LADs, respectively.

**Figure 3:**
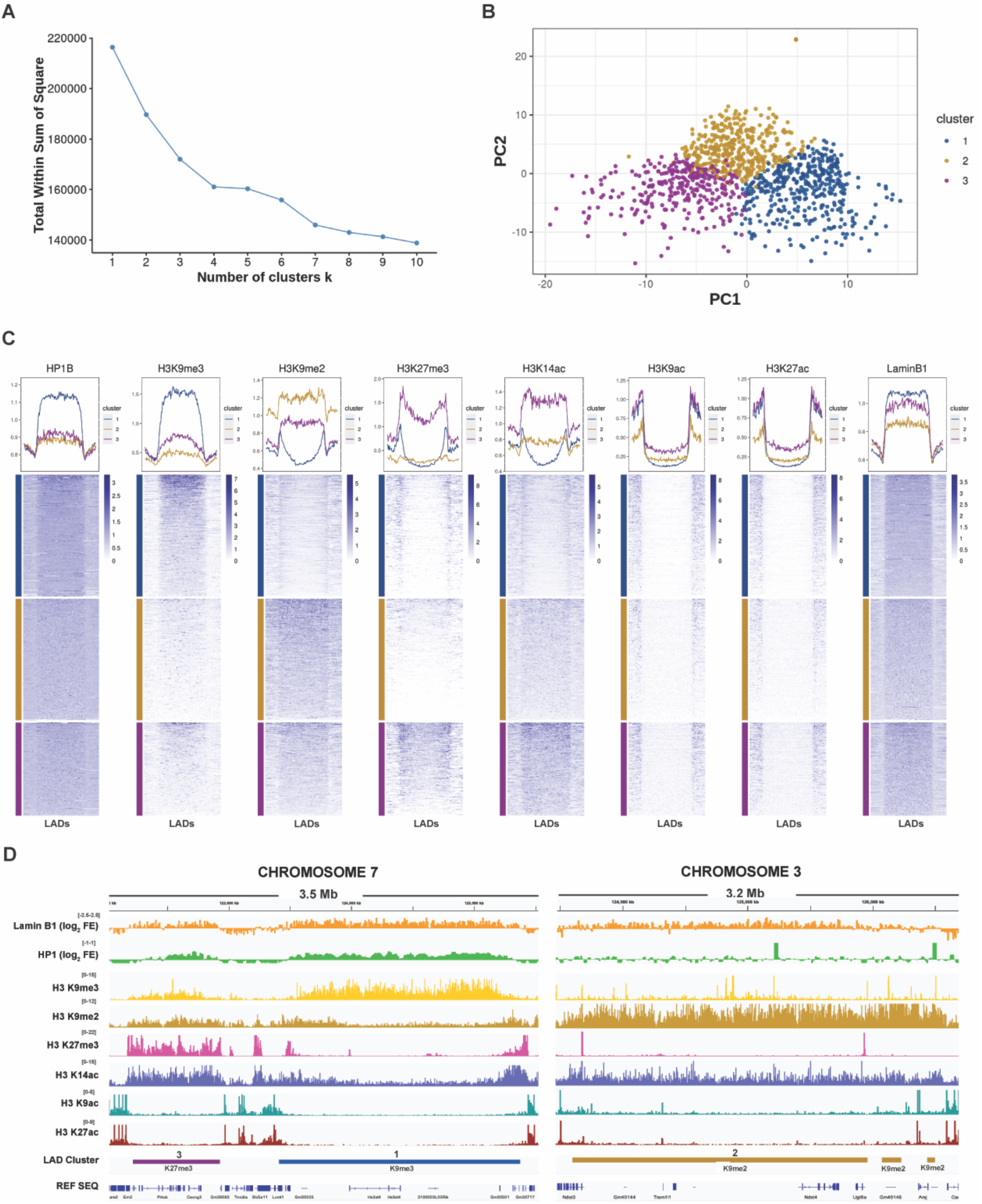
Clustering LADs by chromatin profile. (A) Elbow plot of k-means clustering of LADs. (B) Principle component plot of LAD clusters. The gold dot at the top of the plot consists of about two dozen LADs and is the reason the elbow plot suggests 4 clusters rather than 3. (C) Heatmaps and profiles of feature abundance across LADs, separated into cluster 1 (blue), cluster 2 (gold), and cluster 3 (purple). (D) IGV Browser view illustrating examples from each LAD cluster (from left to right): cluster 3 (K27me3 LADs) in purple, cluster 1 (K9me3 LADs) in blue, cluster 2 (K9me2 LADs) in gold.

Examining chromatin profiles across LADs in each cluster revealed stark differences in both the overall abundance of histone PTMs and their pattern over the LAD (Figure 3C, 3D). As expected, H3K9ac and H3K27ac are highly depleted across the lengths of all 3 LAD clusters. Cluster 1 LADs most closely resemble the canonical definition of constitutive heterochromatin in LADs and the structure of LADs observed in aggregate, featuring a broad domain enriched in HP1β and H3K9me3 across the entire LAD. H3K9me2, H3K27me3, and H3K14ac were depleted throughout the interior of cluster 1 LADs and then peaked near the LAD border with H3K27me3 at the end of the LADs and H3K9me2 and H3K14ac peaking internal to the peak of H3K27me3.

Cluster 2 LADs are distinguished by high levels of H3K9me2 across the entire LAD without the prominent peak at the LAD borders observed in cluster 1 (Figure 3C, 3D). Surprisingly, cluster 2 LADs contain only background levels of H3K9me3 and a very low level of HP1β enrichment. Cluster 2 LADs also contain very low levels of H3K27me3 in the LAD interior with smaller peaks at the borders. H3K14ac is moderately enriched in cluster 2 LADs at a consistent level that is slightly higher than that observed in euchromatic domains. Similar to H3K27me3, the peak of H3K14ac at cluster 2 borders is less distinct than in cluster 1.

Cluster 3 LADs are distinguished by high levels of H3K27me3 across the interior of the LADs with higher levels at the LAD borders (Figure 3C, 3D). They are enriched in H3K14ac, which is present across the LAD at levels significantly higher than seen in most regions of euchromatin. H3K9me2 is present at moderate levels and evenly distributed across the LADs with no discernable peak at the borders. Cluster 3 LADs display moderate levels of both H3K9me3 and HP1β.

All LAD clusters show strong enrichment of Lamin B1, though slight differences could be seen in the degree of Lamin B1 association. Unexpectedly, the lowest Lamin B1 enrichment was seen in Cluster 2, despite several previous studies suggesting that H3K9me2 makes the most significant contribution to lamina association of the histone PTMs studied.

Interestingly, each cluster was characterized by enrichment of a different heterochromatin mark: H3K9me3 for Cluster 1, H3K9me2 for Cluster 2, and H3K27me3 for Cluster 3. To learn which marks contribute most to the differences between clusters, we trained a Random Forest algorithm on our clustering data and looked at which columns were most significant in informing the algorithm’s categorization in the resulting decision tree (Figure S3A). Consistent with our observations, many of the columns contributing most to the differences between clusters belong to H3K9me2, H3K9me3, and H3K27me3, with H3K9me2 and H3K27me3 being particularly significant. H3K14ac and HP1β also make notable contributions, as their relative distributions between clusters resemble those of H3K27me3 and H3K9me3, respectively. Note that for all of these marks, nearly every column driving the differences between clusters is found in the LAD interior (as opposed to the inner or outer border of the LAD), implying that the LAD interior is where clusters differ most. Further supporting the preeminence of H3K9me2, H3K9me3, and H3K27me3 in defining LAD groups, we found that clustering based on only those 3 marks yields quite similar groups, with 916/1128 LADs (81.2%) being assigned to the same clusters even with the reduced set of PTMs (Figure S3B).

Together, these results show that LADs in iMEFs can be divided into three distinct groups which are each primarily characterized by a different heterochromatic PTM, with H3K9me3 predominant in cluster 1, H3K9me2 predominant in cluster 2, and H3K27me3 predominant in cluster 3. This raises the possibility that different regulatory and tethering mechanisms function in each cluster. Given their distinguishing methylation enrichments, we chose to refer to clusters 1, 2, and 3 as K9me3 LADs, K9me2 LADs, and K27me3 LADs, respectively.

### LAD Clusters Are Enriched for Different Genomic Features

Examination of the 3 LAD clusters shows many differences beyond their histone PTM profiles. With a median size of 1.52 Mb, K9me3 LADs are significantly larger than K9me2 LADs and K27me3 LADs, which have median sizes of 0.30 and 0.24 Mb, respectively (Figure 4A). As a result, K9me3 LADs contain the most genes. Taking the LAD size into account, however, shows that the gene density is quite similar between clusters but is highest in K27me3 LADs (Figure 4B). Based on RNA-Seq analysis from these cells, we calculated the total number of reads per Mb in each LAD cluster (Figure 4C). There is no statistical difference in the level of transcription between K9me3 LADs and K27me3 LADs, while there is a slightly lower level of transcription observed in K9me2 LADs. The three LAD clusters also differ in repeat enrichment. K9me3 LADs and K9me2 LADs are highly enriched in LINEs and K27me3 LADs are moderately enriched in SINEs, while LTR enrichment is similar across all groups (Figure 4D).

**Figure 4:**
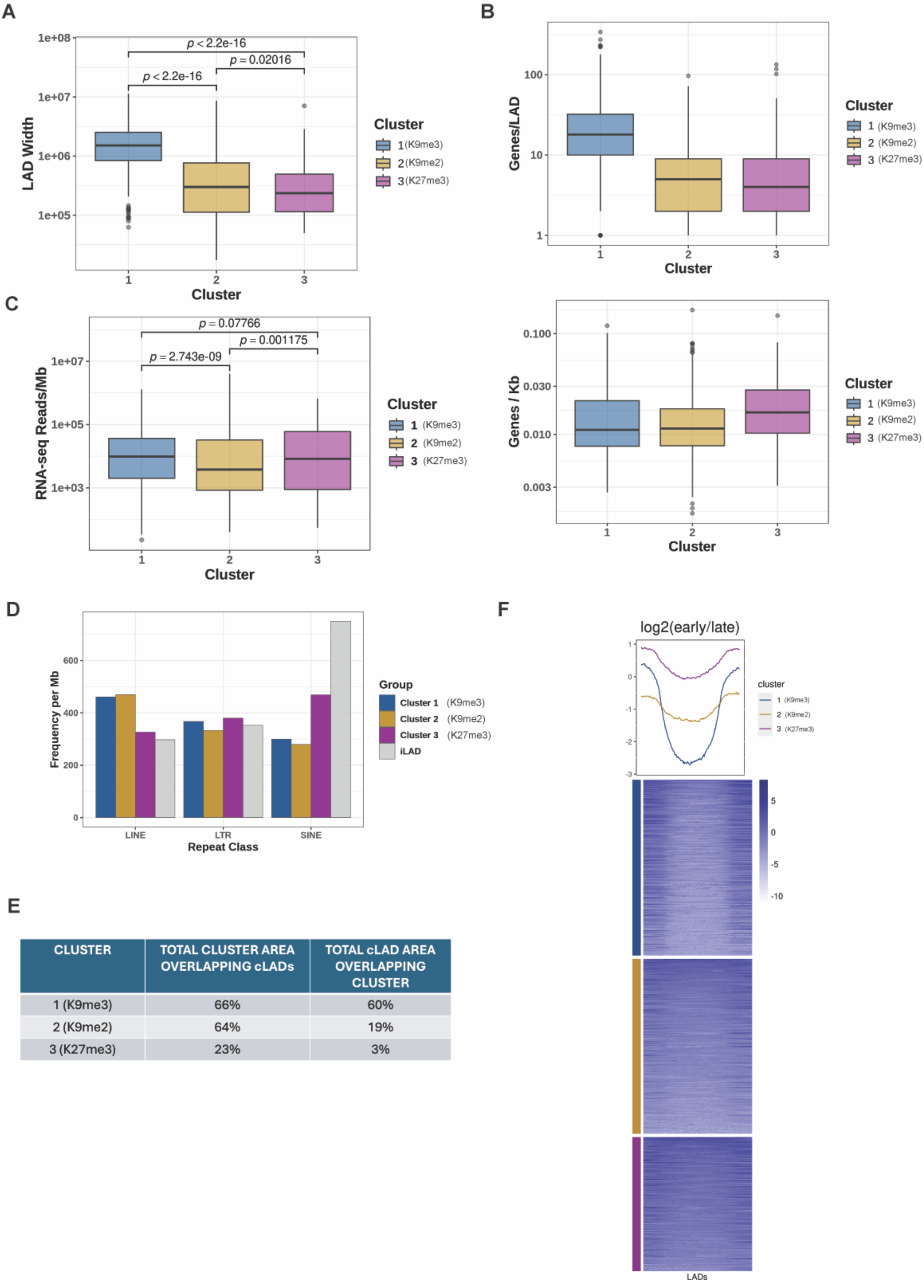
Comparison of K9me3 LADs, K9me2 LADs, and K27me3 LADs. (A) Comparison of LAD sizes (bp) between clusters. P-values determined by Wilcoxon rank-sum test. (B) Gene content in each cluster, represented by total number of genes contained in LADs from each cluster (top) and gene density in each cluster (bottom). (C) Gene expression from LADs of each cluster determined by analysis of RNA-Seq data. P-values determined by Wilcoxon rank-sum test. (D) Abundance of LINE elements, SINE elements, and LTR repeats in each cluster compared to inter-LADs. (E) Table showing overlap between each cluster and cLADs. Note that the cLADs set used came from a previously published DamID map of LADs in several cell types (including MEFs). The total cLAD area overlapping our clusters does not add up to 100% because a small fraction of the LADs detected in those cells were not detected in our LAD mapping which gave rise to the clusters. (F) Heatmap and profile of replication timing across LAD clusters. Values represent log2 early/late signal, so low values indicate late replication timing and higher values represent earlier replication timing.

There is a striking difference in the distribution of cLADs and fLADs between the clusters. Constitutive LADs predominate among K9me3 LADs and K9me2 LADs while the majority of K27me3 LADs are facultative (Figure 4E). This distribution of constitutive and facultative LADs is consistent with the well-characterized involvement of Polycomb silencing in the regulation of developmentally regulated genes(Atlasi and Stunnenberg 2017; Jiang et al. 2024).

LADs are typically associated with late-replicating regions of chromatin, though differences in replication timing between LADs have been observed which correlated with the dynamics of their association with the nuclear lamina during the cell cycle(van Schaik et al. 2020; Gholamalamdari et al. 2024). Using publicly available Repli-seq data in MEFs(Rivera-Mulia et al. 2018), we calculated the log_2_ ratio of early S-phase to late S-phase replication signal and plotted the distribution between clusters. While all three clusters did indeed favor late replication, there were statistically significant differences between clusters. K27me3 LADs, in particular, showed a tendency towards earlier replication, while K9me3 LADs most heavily favored late replication (Figure 4F).

Together, these results suggest that the unique chromatin profiles in each cluster represent underlying biological differences in their function and regulation. K9me3 LADs best resemble the classic definition of a LAD, while K27me3 LADs represent a unique LAD chromatin signature deficient in the H3K9me2/3 normally associated with LADs while strongly enriched in H3K27me3 and H3K14ac. K9me2 LADs appear in many regards to fall between the other two clusters, but also exhibits unique characteristics, such as enrichment of H3K9me2 and lack of H3K9me3.

### chromHMM Confirms Unique Enrichment of States Between Clusters

To further understand and validate the differences observed between LAD clusters, we ran chromHMM with our histone modification, HP1β, and Lamin B1 datasets. ChromHMM uses a Hidden Markov Model to identify underlying chromatin states based on the presence or absence of features of interest (e.g. histone PTMs) across the genome. We ran the model using 10 kb bins and found 8 states to be optimal for capturing a diversity of chromatin states while avoiding redundancy.

The resulting states span the range of active and repressive chromatin (Figure 5A). The model identifies one “euchromatin” state enriched in H3K9ac and H3K27ac (state 8), and one state generally depleted in all histone marks, which appears in various regions across the genome (state 1). The model also identifies one “facultative heterochromatin” state enriched in H3K27me3 (state 7). There are five states primarily populated by H3K9 methylation: two characterized by H3K9me2 (states 5 and 6), two characterized by H3K9me3 (states 2 and 3), and one moderately enriched in both (state 4). Interestingly, Lamin B1 was primarily found in states containing H3K9me3. We were also interested to find that H3K14ac was not found in the euchromatic state 8. Rather, H3K14ac was found in both facultative and constitutive heterochromatin states in association with H3K27me3 and H3K9me2.

**Figure 5:**
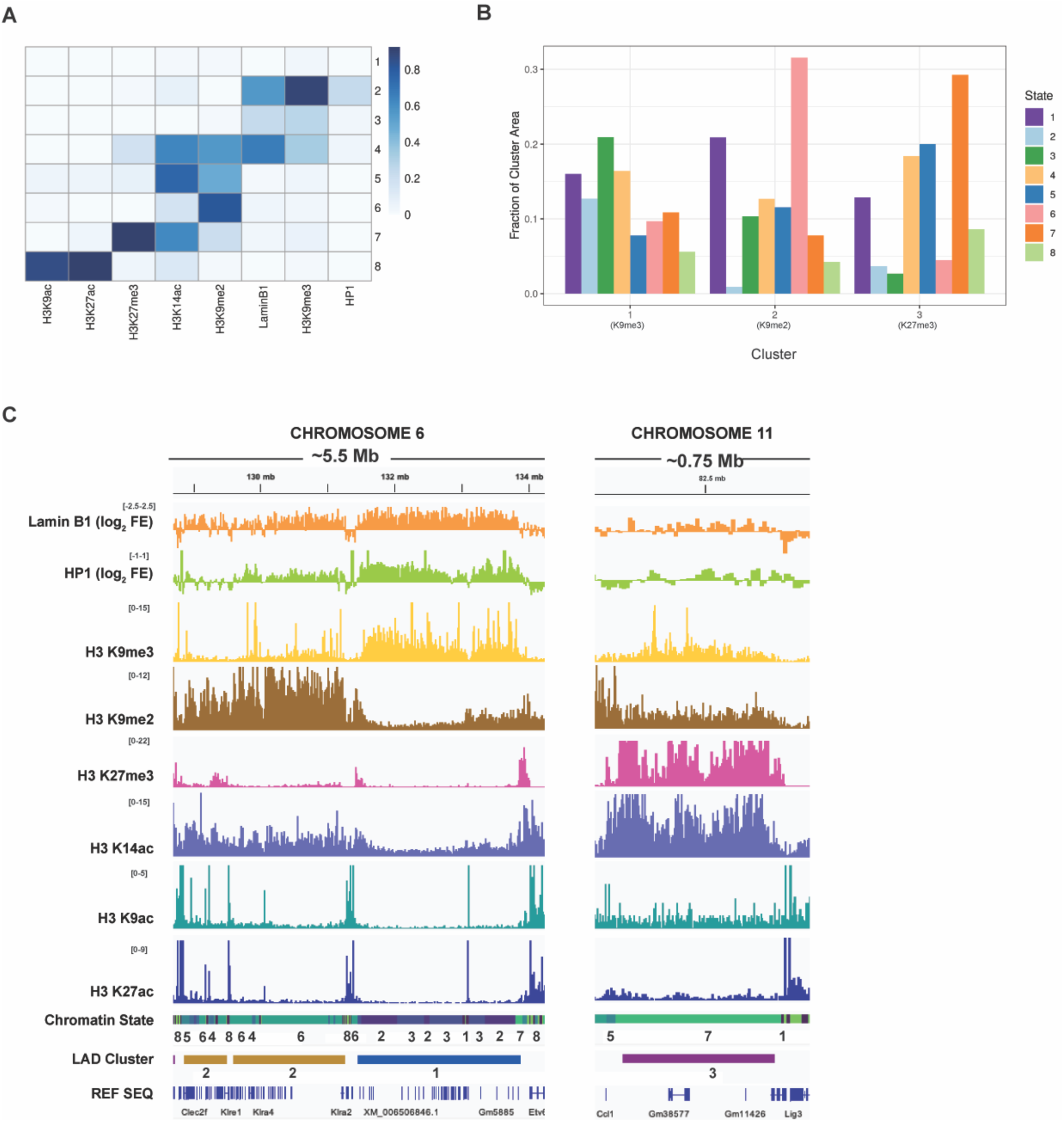
chromHMM analysis and comparison with LAD clusters. (A) Heatmap of feature probabilities in each state. chromHMM was run with 8 states using 10 kb bins. (B) Abundance of each state in cluster 1 (K9me3 LADs), cluster 2 (K9me2 LADs), and cluster 3 (K27me3 LADs). Height of bars represents the number of 10 kb bins within the LAD cluster that corresponded to that state. (C) IGV Browser view showing example LADs from each cluster and the distribution of chromHMM states across the LAD (e.g., enrichment of states 4-6 in K9me2 LADs, states 1-3 in K9me3 LADs, and state 7 in K27me3 LADs).

To compare our observations on the structure of LADs with the states output by chromHMM, we examined the enrichment of states in each LAD cluster (Figure 5B, 5C). As expected, states enriched in K9me3 LADs preferentially contained H3K9me3, the most abundant being states 3 and 4. Also consistent with expectations, the H3K9me2-enriched state 6 predominated in K9me2 LADs and the H3K27me3-enriched state 7 was the most abundant in K27me3 LADs. Inter-LAD regions were largely enriched in the euchromatic state 8 and the more variable state 1.

### LAD clusters contain distinct states at their borders

Examination of states at LAD borders confirms differences in the border structure of different LAD clusters (Figure 6). Moving from outside a LAD across the border into a LAD, K9me3 LADs display a clear transition from state 8 to more “LAD Border” states enriched in H3K27me3/H3K14ac, to constitutive heterochromatin inside the LAD (Figure 6A, 6B). However, this transition is altered in K9me2 LADs and K27me3 LADs, which retain a high abundance of H3K9me2 and H3K27me3, respectively across both the LAD interior and borders. K9me2 LADs have little to no enrichment of H3K27me3 or H3K14ac at borders, appearing to lack this usual LAD barrier (Figure 6A, 6C). H3K27me3 and H3K14ac remain enriched at K27me3 LAD borders but are also enriched in LAD interiors (Figure 6A, 6D).

**Figure 6:**
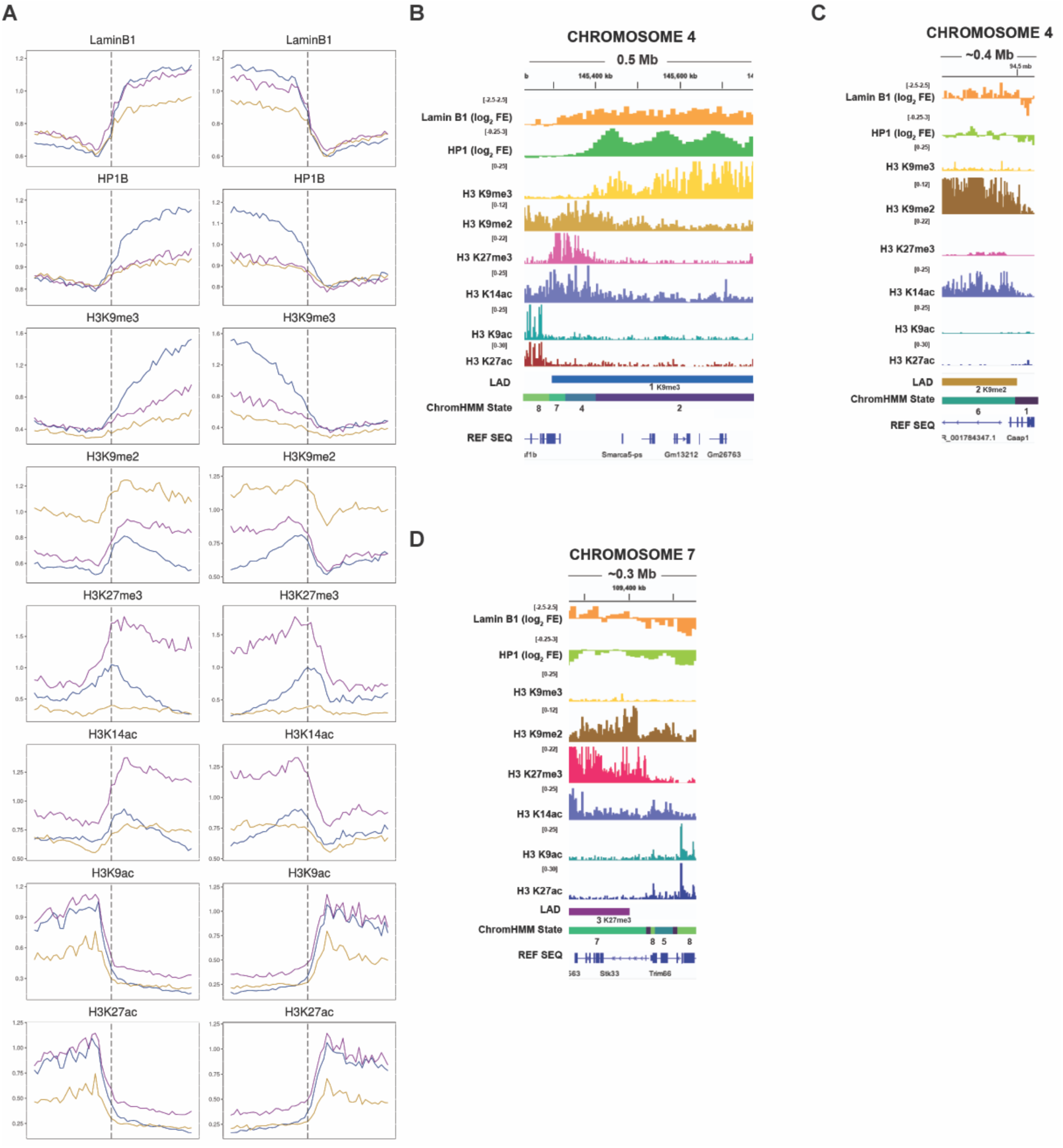
LAD border profiles vary between clusters. (A) Profiles of all features around LAD borders. Boxes show a 1 Mb window centered around the LAD borders (vertical dotted line). Note that the profiles extend 500 kb inside the LAD border, longer than the full length of some LADs, and thus include the opposite LAD border. This is especially true for K9me2 and K9me3 LADs, most of which are significantly less than 500 kb long. Thus, the apparent continual decrease of H3K27me3 and H3K14ac inside LADs, which appears at odds with the scaled profiles in Figure 3, is likely a result of increasing inter-LAD regions being included as the profile is traced away from the LAD border and beyond the opposite border of these short LADs. The apparent enrichment of Lamin B1 at the borders of K27me3 LADs is like a eresulkt of the same phenomenon. (B) IGV Browser example of epigenetic features and chromHMM state transitions across a cluster 1 LAD (K9me3 LAD) boundary. (C) IGV Browser example of epigenetic features and chromHMM state transitions across a cluster 2 LAD (K9me2 LAD) boundary. (D) IGV Browser example of epigenetic features and chromHMM state transitions across a cluster 3 LAD (K27me3 LAD) boundary.

### Regression analysis suggests HP1-independent tethering mechanisms and differences in cluster regulation

Many mechanisms contribute to the localization of chromatin to the nuclear periphery and they remain poorly understood. Most involve interactions of peripheral proteins with H3K9me2/3, often mediated by the H3K9me2/3-binding protein HP1, which can bind lamina-associated factors such as LBR and PRR14. To learn which factors or mechanisms were responsible for mediating chromatin-lamina interactions in C57/bl6 MEFs and whether these differed between clusters, we performed pairwise regression analysis between the net abundance of various histone PTMs, HP1β, and Lamin B1 in LADs. As expected, we observed a consistent positive correlation between the abundance of HP1β and its substrate, H3K9me3 when looking at total LADs and this positive correlation was observed for each LAD cluster (Figure S4A). A strong positive correlation was also seen between HP1β and Lamin B1 and between H3K9me3 and Lamin B1 with all LADs (Figure S4B, S4C). These relationships are preserved in each of the individual clusters, though the strength of the correlation is significantly reduced. Unexpectedly, the relationship between H3K9me3/HP1β and Lamin B1 was weakest in the group where they were most enriched, K9me3 LADs. K9me3 LADs displayed a similar correlation between H3K9me2 and Lamin B1, and an even stronger correlation between H3K27me3 and Lamin B1, despite being depleted in both H3K9me2 and H3K27me3 (Figure S4D, S4E).

Similarly, Lamin B1 increased with the abundance of H3K9me3 and H3K27me3 in K9me2 LADs but showed no correlation with H3K9me2 (Figure S4C, S4D, S4E). K27me3 LADs showed no correlation between H3K27me3 and Lamin B1 and only weak correlation between H3K9me2 and Lamin B1 (Figure S4D, S4E). H3K14ac generally correlated with other features in a similar manner to H3K27me3, consistent with their similar patterns of enrichment across LAD clusters (Figures 3C, 6A, S4F, S4G). Notably, PTMs that correlated positively with Lamin B1 did not necessarily correlate with HP1β and vice-versa, suggesting in at least some cases HP1β was not mediating interactions with the nuclear lamina.

## Discussion

Prior studies have pointed to structural and functional differences within LADs. Our analysis of LADs clustered by Lamin B1, HP1β, H3K9me3, H3K9me2, H3K27me3, H3K14ac, H3K9ac, and H3K14ac demonstrated that LADs in MEFs fall into 3 distinct subtypes with significant differences in their epigenomic and genomic content. Each of the 3 clusters we identified has a distinct and prominent marker of heterochromatin. Cluster 1 is characterized by high levels of H3K9me3 (and is therefore referred to as K9me3 LADs), cluster 2 has high levels of H3K9me2 (called K9me2 LADs) and cluster 3 contains high levels of H3K237me3 (called K27me3 LADs). This study has the advantage of having mapped these features in cell lines derived from the same colony of C57BL/6 mice, avoiding possible discrepancies that could arise when using public datasets generated by different labs in their own MEF cell lines.

Our approach of scaling LADs to a uniform size and dividing them into bins prior to K-means clustering provides several advantages for grouping LADs. First, it gives all LADs equal weight so that large LADs do not dominate during clustering and mask differences found in small LADs. Second, it allows different patterns of the same mark across LADs to be distinguished (e.g., H3K14ac enriched uniformly across LADs in K9me2 LADs vs. specifically at borders in K9me3 LADs). Dividing LADs into bins allows differentiation of LADs which vary in the distribution of marks across the LAD, even when the overall abundance is similar.

K9me3 LADs and K9me2 LADs most resemble the classical definition of LADs: large, gene-poor, and enriched in LINE elements and methylation of H3K9. Surprisingly, however, few LADs were enriched in both H3K9me2 and H3K9me3. K9me3 LADs were depleted in H3K9me2 everywhere except at the borders, and K9me2 LADs contained only very low enrichment of H3K9me3 or HP1β. This suggests that H3K9me2 and H3K9me3 rarely coincide in LADs, perhaps because they are mutually exclusive on the same histone. In some LADs, therefore, the higher methylation state predominates, while in others, H3K9me2 is maintained and conversion to H3K9me3 prevented. In fact, regression analysis shows that there is a significant negative correlation between the levels of H3K9me2 and H3K9me3 in K9me2 LADs and K9me3 LADs (data not shown). We hypothesize that H3K9 methylation is maintained primarily by Suv39H1/2 in the former and G9a/GLP in the latter, as these primarily catalyze H3K9 trimethylation and demethylation, respectively(Rice et al. 2003; Padeken et al. 2022). This may help explain why both sets of methyltransferases are important for lamina association(Bian et al. 2013; van Steensel and Belmont 2017).

Previous studies have affirmed the existence of small LADs with facultative heterochromatin character, but these are not well studied in comparison with large, canonical LADs(Tran et al. 2021; Alagna et al. 2023; Gholamalamdari et al. 2024). Though H3K27me3 has been proposed to play a role in attaching chromatin to the nuclear periphery, it is generally thought to be restricted to LAD borders. This conception may be partly due to studying LADs as a single, uniform group. By clustering LADs, we uncovered a group of LADs enriched in H3K27me3 across the entire LAD body, confirming that some LADs are indeed characterized by facultative heterochromatin. These LADs are smaller and more enriched in genes and SINE elements than LADs from other clusters. They constitute approximately a third of LADs in iMEFs, but due to their small size they compose only a small fraction of the total lamina-associated genome, which may explain why they are not apparent when characterizing total LAD chromatin. One study has indicated that these LADs are more variable and interact with the nuclear periphery for only part of the cell cycle, which is consistent with their facultative heterochromatin character(Tran et al. 2021).

Intriguingly, a recent study in human cells supported similar LAD divisions, noting four different LAD clusters which were H3K9me3-enriched, H3K27me3-enriched, H3K9me2-enriched, and PTM- depleted, respectively(Gholamalamdari et al. 2024). This study had clustered LADs based on their degree of association with the nuclear lamina and with nuclear speckles, thereby suggesting that these epigenetically distinct groups of LADs are subject to different regulatory mechanisms and behaviors across the cell cycle and development.

Surprisingly, we saw only minor differences in the degree of lamina association between clusters. Given the emphasis of previous studies on the importance of H3K9me2 in lamina association, we expected K9me2 LADs to have the most interaction with the nuclear lamina. Instead, it had the least (albeit by a small margin). However, this finding is not at odds with previous studies as it may at first seem. The relative expression levels of different lamins and lamin-associated proteins change across development and vary between tissues, suggesting that different tethering mechanisms predominate in different cell types. While this study was performed in MEFs, most studies emphasizing the exclusive role of H3K9me2 have been performed in other cell types(Kind et al. 2013; Poleshko et al. 2017). One group found that H3K9me3 levels tended to correlate positively with changes in lamina association in mesenchymal cell lineages (including fibroblasts) and negatively in others(Das et al. 2023). It is therefore possible that differential expression or post-translational modification of lamins or other proteins involved in mediating chromatin-lamina interactions cause different histone PTM-dependent mechanisms to predominate in different cell types. Consistent with this, it has been demonstrated that H3K9me2 is neither necessary nor suYicient^37^ for association of chromatin with the nuclear periphery in all cell types(Towbin et al. 2012; Bian et al. 2013; Harr et al. 2015). One study examining the role of H3K27me3, H3K9me2, and H3K9me3 in chromatin-lamina tethering in MEFs concluded that all 3 are required for proper chromatin-lamina interactions(Harr et al. 2015). Another complemented these results by finding that H3K9me2 and H3K9me3 are both enriched at the nuclear periphery in MEFs but are anticorrelated, suggesting that they mark different regions governed by different peripheral targeting mechanisms(Bian et al. 2013).

Consistent with distinct roles and tethering mechanisms at work between clusters, our regression analysis showed differences between clusters regarding which features positively correlated with Lamin B1. Surprisingly, the heterochromatin mark which was most enriched in each cluster was also the least correlated with Lamin B1 signal. Three explanations for this may be proposed. First, these clusters may be saturated with their respective heterochromatin marks, preventing them from covarying with Lamin B1 association. Second, other mechanisms may be at work promoting and/or antagonizing lamina association in these clusters independently of the predominant heterochromatin mark. Third, the predominant heterochromatin mark may be involved simply in silencing chromatin without regulating association with the nuclear lamina. While each cluster is highly enriched in its respective mark, none appear to be constitutively associated with the nuclear periphery, since their Lamin B1 association varies significantly. Thus, the first option is unsatisfactory on its own. However, some combination of options 1 and 2 is possible and may be preferable to the third option, which would raise the question of how any of these LADs are then tethered and why heterochromatin marks with the ability to promote lamina-association do not do so in the areas where they are most highly enriched.

The cell has numerous mechanisms to establish and maintain borders between euchromatin and heterochromatin. This is underscored by our observations of the complexity and uniqueness of LAD borders. In addition to the well-characterized enrichment of H3K27me3, previous studies have identified enrichment of factors such as CTCF, macro-H2A, and gene promoters in the vicinity of LAD borders(Guelen et al. 2008; Fu et al. 2015; Harr et al. 2015; van Schaik et al. 2022). We have now identified a new feature of LAD borders, H3K14ac. H3K14ac is generally associated with transcriptional activation and has received relatively little attention compared to many other acetylation marks(Wang et al. 2008; Karmodiya et al. 2012; Regadas et al. 2021). Previous studies have shown that it is a crucial regulator of heterochromatin maintenance in fission yeast, marks a subset of promoters and enhancers in higher eukaryotes, and is essential for mammalian development(Kueh et al. 2011; Reddy et al. 2011; Karmodiya et al. 2012; Stirpe et al. 2021; Kueh et al. 2023; Seman et al. 2023). Our results show that H3K14ac accumulates just inside LAD borders and gradually decreases into the middle of the LAD. This stands in stark contrast to H3K9ac or H3K27ac which are highly depleted across LADs and enriched outside LADs, and suggests a role for H3K14ac in regulating LAD borders. Many LADs (most K9me2 LADs and K27me3 LADs) show moderate to high enrichment of H3K14ac across the entire LAD body as well as the borders. These findings raise many interesting questions regarding the role of H3K14ac in regulation of chromatin-lamina interactions.

We also found a moderate enrichment of H3K9me2 inside LAD borders, particularly in K9me3 LADs, which were otherwise depleted in H3K9me2. While H3K9me2 is well-studied in LADs, it has generally been seen across LAD bodies, pointing to another potential difference between cell-types and LAD clusters. Interestingly, H3K14ac and H3K9me2 at LAD borders do not align with the well-characterized peaks of H3K27me3 at LAD borders, and are instead shifted inward. The purpose for this localization is not clear, but may suggest that these marks reenforce the border by forming a second barrier against the euchromatin outside LADs. Combined with H3K27me3, H3K14ac, and H3K9me2, these findings underscore the complexity and regulatory importance of LAD border regions.

The results of this study contribute to a growing understanding of the internal distinctions between LADs and reveal significant features shaping these differences and contributing to LAD function and regulation. They also highlight how LAD regulation may differ across different cell types, underscoring the complexity with which chromatin-lamina interactions and the mechanisms governing those interactions are coordinated and adjusted through development and differentiation.

## Methods

### Cell Culture

Mouse embryonic fibroblasts harvested and immortalized as previously described were cultured at 37 °C with 5% CO_2_(Nagarajan et al. 2013). Cells were grown in Dulbecco’s Modified Eagle’s High Glucose Medium supplemented with 10% FBS, 1% glutamine, and 1% Penn-Strep and passaged approximately every 2-3 days. Cells were harvested for experiments using 0.5% Trypsin and resuspended in fresh media for counting.

### CUT&Tag

CUT&Tag was performed according to the EpiCypher® CUTANA™ Direct-to-PCR CUT&Tag Protocol, with a light crosslinking step adapted from Kaya-Okur et al. (2020)(Kaya-Okur et al. 2020). In short, 100,00 MEFs were harvested and rinsed with PBS, then incubated in Nuclear Extraction Buffer for 10 minutes on ice. Nuclei were spun down and resuspended in PBS, then fixed by adding formaldehyde to a concentration of 0.1% and incubating 2 min at room temperature. The formaldehyde was quenched with two molar equivalents of glycine and cells were spun down, resuspended in Nuclear Extraction Buffer, then added to 10 uL activated Concavalin A beads and incubated for 10 minutes at room temperature for nuclei to bind to the beads. The nuclei/beads were separated with a magnet to allow removal of the supernatant, suspended in Antibody 150 Buffer with the appropriate dilution of primary antibody and incubated overnight at 4 °C. The next day, the primary antibody was replaced with secondary antibody and nuclei were incubated for 30 min at room temperature. Nuclei were rinsed twice with Digitonin 150 Buffer and incubated for 1 hour with pAG-Tn5 fusion transposase, followed by two rinses with Digitonin 300 Buffer. They were then incubated at 37 °C for 1 hour with 10 mM MgCl_2_ to activate the Tn5 Transposase. Tagmentation was stopped by rinsing with TAPS Buffer and samples were incubated in SDS Release Buffer at 58 °C for 1 hour to lyse nuclei and release the DNA. SDS was quenched in 0.5% TritonX-100. DNA libraries were amplified using CUTANA High-Fidelity 2X PCR Master Mix™ with Universal i5 primer and the appropriate barcoded i7 primer, and DNA was extracted using 1.3X AMPure beads. Libraries were sequenced with paired-end Illumina sequencing. All CUT&Tag experiments were performed in triplicate using three independently derived cell lines from the same mouse colony.

### CUT&RUN

CUT&RUN was performed according to the EpiCypher® CUT&RUN Protocol. 500,000 iMEFs were harvested, rinsed twice with wash Buffer, bound to activated Concavalin A beads, and incubated overnight with primary antibody at the appropriate dilution in Antibody Buffer. The next day, samples were rinsed twice with Digitonin Buffer and incubated for 10 minutes with pAG-MNase. There were then rinsed twice more with Digitonin Buffer and incubated in 2 mM CaCl_2_ for 2 hours at 4 °C. Samples were quenched with 33 uL STOP Buffer and incubated 10 min at 37 °C to release fragmented DNA, then beads were separated out of solution on a magnetic rack and the supernatant was transferred to new tubes. Libraries were prepared for next generation sequencing using the CUTANA CUT&RUN Library Prep Kit. Lamin B1 CUT&RUN was performed in duplicate. CUT&RUN for H3K9me2 and H3K14ac to validate CUT&Tag results was performed in single replicate and a nuclei extraction step was inserted prior to bead binding as in CUT&Tag.

### ChIP-seq and RNA-Seq

HP1β ChIP-seq was performed as previously described for H3K9me2 and H3K9me3**(Popova et al. 2021).** RNA-seq was performed in as described (Nagarajan et al. 2019).

### Data pre-processing and LAD calling

Before alignment, raw CUT&RUN data was trimmed using TrimGalore with the arguments *-q 20 -- stringency 3 --trim-n*. CUT&Tag and CUT&RUN data were aligned to the mm10 genome using bowtie2 with parameters *--end-to-end --very-sensitive --no-unal --no-mixed --no-discordant -- phred33 -I 10 -X 700*. HP1β ChIP-seq data was aligned to the mm10 and hg19 genomes using BWA mem with flag *-M*. Data was filtered to remove reads mapping to mitochondrial DNA or unlocalized or unplaced contigs. Individual replicates were merged and converted to bedGraph or bigwig format for viewing on IGV and for downstream analysis.

For LAD calling, Lamin B1 and IgG CUT&RUN data were normalized to the same library size using samtools view, filtered to remove reads overlapping the ENCODE ChIP-seq blacklist, and replicates were merged. LADs were called using the HMMt package in R with a modified version of the HMMtBroadPeak function which used a sliding window to smooth the data by averaging a bin’s signal with the two adjacent bins. LADs were first called by the modified function using 30 kb bins. The ends of the LADs were then adjusted to increase the resolution by running the normal HMMtBroadPeak function with 5 kb bins and using the output to trim or extend LAD ends depending on whether the overlapping 5 kb bin was LAD positive (Supplementary File S1).

### Data clustering

For clustering, histone PTM, HP1β, and Lamin B1 bed files were filtered to remove reads overlapping the ENCODE ChIP-seq blacklist and converted to bigWig format. We then used deepTools computeMatrix to scale all LADs to a size of 1 Mb and summarize bigWig signal into 50 kb bins across the LAD, including 100 kb outside the LAD border. The data was read into R and filtered to remove chrX and chrY, and outliers were conservatively removed by going through each column of the matrix and replacing all values more than 1.5 times the interdecile range away from the median with the value of the 1^st^ or 9^th^ decile for low or high outliers, respectively. Each column was then scaled to a mean of 0 and standard deviation of 1 and the data was clustered using the kmeans function in R. The factoextra package was used to create elbow and silhouette plots to determine the optimal number of clusters. Hierarchical clustering was performed using the hclust function in R with Ward’s Linkeage. To learn which marks defined the differences between clusters, we used the randomForest package and ran the algorithm on the entire dataset since our intention was simply to train the model on our data, not make predictions.

Profiles and heatmaps of the resulting clusters were created with ggplot2 and pheatmap using a matrix generated and filtered in the same way as the one used for clustering but with 10 kb bins for better resolution and with 300 kb included beyond each LAD border.

### chromHMM

chromHMM 1.24 was run on bam files for Lamin B1, HP1β, H3K9me3, H3K9me2, H3K27me3, H3K14ac, H3K9ac, and H3K27ac. We used a 10 kb bins size for binarizing bam files, then ran LearnModel with 8 states and a 10 kb bin size, specifying the mm10 genome. The dense bed file was used for further analysis after first filtering to remove all bins which overlapped gaps of unknown nucleotides more than 5 kb long.

The enrichment of states in each cluster was calculated by adding up the number of bins for each state which overlapped LADs of that cluster.

### Cluster analysis

Data analysis and visualization were performed in R. Gene annotations for the RNA-seq and to calculate gene enrichment in each cluster came from GENCODE vM25. RNA-seq data was analyzed using the nf-core rnaseq pipeline version 3.13.12 with Star and Salmon and aligned bam files were loaded into R to determine the number of reads overlapping each LAD. The matrix to create the profile and heatmap of replication timing was created by generating matrices of the repli-seq signal from early and late S-phase using deepTools as with the other profiles and heatmaps, then calculating the log_2_ of the early replication signal divided by the late replication signal for each bin. Repeat enrichment was obtained from RepeatMasker through the UCSC table browser and repeat enrichment in each cluster was calculated by counting the number of repeats of each category that overlapped LADs in that cluster.

Boxplots, barplots, and Venn diagrams were made with ggplot2 and p-values for the boxplots were calculated by Wilcoxon rank sum test using the ggsignif package. Heatmaps were made with the pheatmap package.

### Linear Regression

To perform linear regression, the total CUT&Tag/CUT&RUN/ChIP-seq signal for the mark of interest in each LAD was summed up using bedtools. The data was loaded into R and filtered to remove LADs in sex chromosomes, and the signal in each LAD was divided by the LAD size to get the average abundance of the mark across the LAD and avoid skewing values by LAD size. The data was log_2_-transformed to achieve a more normal distribution and all points were dropped which fell more than 3 standard deviations from the mean in either of the two marks being compared. The number of LADs being dropped as outliers ranged from 7 to 24 depending on the marks. Regression was performed using the linear model function in base R and the p-value of the slope was reported on the plot.

### Data sources

Gene annotations for GRCm38.p6 were downloaded in GTF format from GENCODE release M25 (https://www.gencodegenes.org/mouse/release_M25.html) and chromosomes were renamed to UCSC format for consistency with the rest of our analysis.

Repli-seq data came from 4D Nucleome (data files <u>4DNFI9LSI1RE</u> and <u>4DNFID3GEVFS</u>)(Rivera-Mulia et al. 2018).

Repetitive Element Annotations were obtained from RepeatMasker through UCSC table browser DamID maps of LADs and cLADs in MEFs came from Gene Expression Omnibus (GSE17051)^4,52^ and coordinates were converted from the mm9 to mm10 genome using UCSC LiftOver(Peric-Hupkes et al. 2010; Meuleman et al. 2013).

### Antibodies

HP1β: 8676 (Cell Signaling)

H3K9me3: ab8898 (abcam)

H3K9me2: ab1220 (abcam) for CUT&Tag and main analysis

H3K9me2: 39041 (Active Motif) for CUT&RUN validation

H3K27me3: 9733 (Cell Signaling)

H3K14ac: ab52946 (abcam) for CUT&Tag and main analysis

H3K14ac: MA5-32814 (ThermoFisher) for CUT&RUN validation

H3K27ac: 8173 (Cell Signaling)

H3K9ac: 9649 (Cell Signaling)

Lamin B1: ab16048 (abcam)

IgG: 13-0042 (Epicypher)

## Data Access

The datasets generated in the current study are available at the GEO database. CUT&RUN, Cut&Tag, ChIP-Seq, and RNA-Seq datasets have the accession numbers: GSE281928, GSE281927, GSE281926, and GSE281925, respectively.

## Acknowledgements

The authors would like to thank Dr. Michael Freitas for meany helpful discussions. This work was supported by grant R01 GM144601 from the National Institutes of Health.

